# Comparative study of curvature sensing mediated by F-BAR domain and an intrinsically disordered region of FBP17

**DOI:** 10.1101/2020.07.30.230037

**Authors:** Maohan Su, Yinyin Zhuang, Xinwen Miao, Yongpeng Zeng, Weibo Gao, Wenting Zhao, Min Wu

**Author notes:** Equal contribution (Maohan Su, Yinyin Zhuang).

## Abstract

Membrane curvature has emerged as an intriguing physical organization principle underlying biological signaling and membrane trafficking. FBP17 of the CIP4/FBP17/Toca-1 F-BAR family is unique in the BAR family because its structurally folded F-BAR domain does not contain any hydrophobic motifs that insert into lipid bilayer. While it has been widely assumed so, whether the banana-shaped F-BAR domain alone can sense curvature has never been experimentally demonstrated. Using a nanopillar-supported lipid bilayer system, we found that the F-BAR domain of FBP17 displayed minimal curvature sensing *in vitro*. We further identified an alternatively spliced intrinsically disordered region (IDR) of FBP17 next to its F-BAR domain that is conserved in sequence across species. The IDR senses membrane curvature and its sensing ability greatly exceeds that of F-BAR domain alone. In living cells, presence of the IDR domain changed the dynamics of FBP17 recruitment in a curvature-coupled cortical wave system. Collectively, we propose that FBP17 does sense curvature but contrary to the common belief, its curvature sensing capability largely originates from its disordered region, not F-BAR domain itself.

## Introduction

Membrane curvature in cells needs to be dynamically generated and sensed by peripheral membrane proteins in biological processes involving membrane deformation. In principle, curvature-sensing proteins should be more abundant than generators because some of the sensors that are too flexible to bend the membrane may fail to generate curvature (Zimmerberg and Kozlov, 2006). In reality, numerous proteins have been reported to deform cellular membranes but much fewer proteins have been shown to sense membrane curvature (Antonny, 2011). While curvature generation and sensing have been proposed as behaviors by the same protein at different concentration regimes (Simunovic et al., 2018; Suetsugu and Gautreau, 2012), whether proteins generating curvature are by default curvature sensors are questioned (Madsen et al., 2010).

Multiple mechanisms have been proposed to mediate curvature sensing, including scaffolding and hydrophobic membrane insertion (Antonny, 2006; Qualmann et al., 2011). The BAR (Bin/Amphiphysin/Rvs) domain superfamily consists of proteins that are known scaffolders. They share in common banana-shaped domains that can bind to membrane via its concave face (Farsad and Camilli, 2003; Kozlov et al., 2014; McMahon and Gallop, 2005). Because of the shape of BAR domains (Frost et al., 2009; Mim et al., 2012; Peter et al., 2004), it was intuitively assumed both curvature generators and curvature sensors. However, many BAR proteins contain amphipathic helices in addition to the crescent shape. Whether a scaffolding mechanism alone can sense curvature remains controversial (Bhatia et al., 2010; Simunovic et al., 2015). Surface-based Single Liposome with Curvatures Assay (SLiC) showed that the N-terminal amphipathic helices without BAR domains can sense curvature while BAR domains without amphipathic helices cannot (Bhatia et al., 2009). On the other hand, in the tube-pulling assay, BAR domains devoid of these helices were still highly curvature-sensitive (Simunovic et al., 2016). One caveat of the tube-pulling assay is that membrane curvature was modulated by changing aspirating pressure, hence curvature is intimately connected with membrane tension and the effect of either factor alone is difficult to isolate.

F-BAR proteins of FBP17/CIP4/Toca sub-family are intriguing examples to study the contribution of scaffolding in curvature sensing. They naturally lack the amphipathic helices or the hydrophobic insertion region present in other BAR domains (Qualmann et al., 2011). Previous experimental evidences on the curvature sensing of the F-BAR domain in this sub-family are lacking. When liposome binding experiments were performed, F-BAR of FBP17 or Toca1 was shown to prefer larger liposomes at 1000 nm diameter rather than smaller ones of 100 nm (Takano et al., 2008). In addition to FBP17/CIP4/Toca family, F-BAR domains from Cdc15 and FCHo2 show no curvature preference (Henne et al., 2007; McDonald et al., 2015). In comparison, using similar assays ENTH domain of Epsin1 could distinguish liposome sizes between 50 to 800 nm (Peter et al., 2004) and F-BAR domain of syndapin1 showed increasing binding towards smaller liposomes (<100 nm diameter) (Ramesh et al., 2013). However, both epsin and syndapin1 contains hydrophobic insertion motifs that could potentially mediate their curvature sensing, further raising the possibility that scaffolding mechanism may not be necessary for curvature-sensing.

In this work, we employed two experimental approaches to address whether the F-BAR domain of FBP17 senses curvature. We used the dynamic pattern formation of mast cells to investigate protein dynamics at the time scale of seconds. We showed that the phase of the F-BAR and BAR protein recruitment to the travelling waves is primarily governed by their membrane binding domains and can be predicted from their curvature preference. We found that the phases of two FBP17 isoforms consistently differ, and the differential spliced intrinsically disordered region (IDR) modulates F-BAR protein’s membrane binding and curvature sensing preferences. Dissecting the relationship between curvature generation and sensing is challenging in living cells because those two processes are likely to be coupled (Suetsugu et al., 2014). We then employed a newly developed nanobar supported lipid-bilayer system to test curvature sensing *in vitro* (Zhao et al., 2017). Using purified proteins, we showed that the F-BAR domain of FBP17 displayed weak but positive curvature sensitivity. However, most of the curvature sensing effects appear to originate from a previously uncharacterized IDR following the F-BAR domain.

## Results

### Phases of protein recruitment to the travelling waves correlate with curvature progression

We previously showed that wave formation in cells is coupled with changes in membrane curvature, and our mechanochemical feedback model predicted that curvature-sensing is the key for fast wave propagation (Wu et al., 2013). Since the changes in membrane curvature must be continuous, we set out to test to what extent the timing of the recruitment of curvature generating proteins, i.e. the phase of the protein in waves, could correlate with their curvature preferences *in vitro*. We first imaged FBP17 (the commonly used short isoform corresponding to rat Rapostlin) and N-BAR domain only (a.a. 1-247) from Endophilin-1 (N-BAR^endo^) (**Fig. 1a**), which has the ability to induce membrane tubules with higher curvature (Mim et al., 2012) than the F-BAR domain from FBP17 (Frost et al., 2008). By co-overexpressing them in RBL cells, we found that N-BAR^endo^ could assemble as dynamic puncta in the form of travelling waves together with FBP17 (**Fig. 1b, c**), and its phase lagged FBP17 for about 5 s (**Fig. 1d**). N-BAR^endo^ was in the same phase as full-length endophilin (**Supp Fig. 1a**). When we replaced the F-BAR domain from FBP17 with N-BAR^endo^, this chimeric protein appeared in waves at the same phase of full-length endophilin but not that of FBP17 (**Supp Fig. 1b**). These data suggest that the timing of endophilin recruitment is determined by its membrane-interacting N-BAR domains and can be modulated by changing its membrane interaction. We also tested the N-BAR domain (a.a. 1-236) from Amphiphysin-1 (N- BAR^amph^), which shares similar intrinsic curvature with N-BAR^endo^ (Mim et al., 2012; Peter et al., 2004). N-BAR^amph^ also formed dynamic puncta as waves and its phase lagged FBP17 by 5 s (**Supp Fig. 1c**), indicating that this phase shift may be dictated by common property of N-BARs.

**Figure 1.**
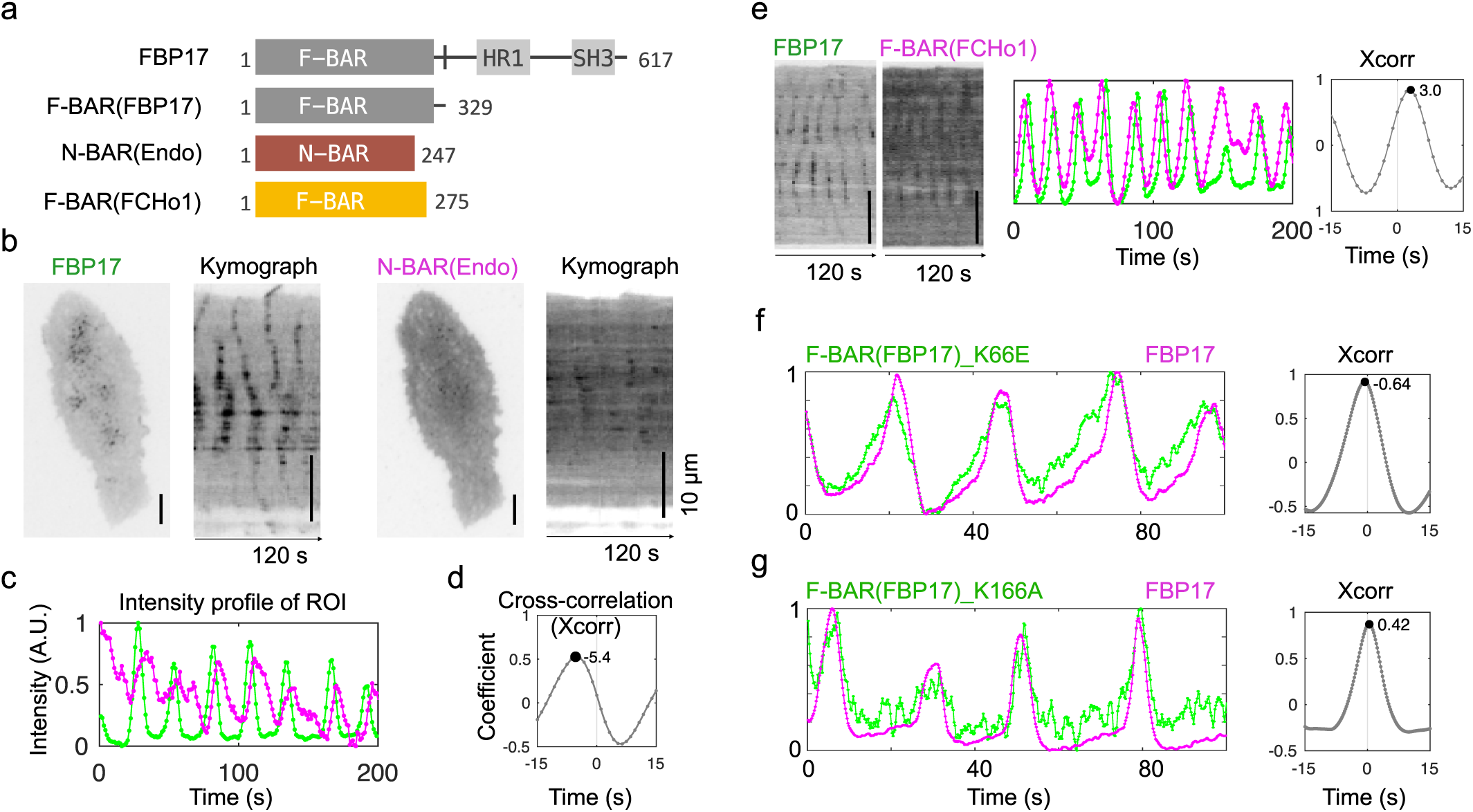
Dynamic recruitment of FBP17, F-BAR and N-BAR to traveling waves. **a**. Diagram of constructs used: FBP17, F-BAR domain from FBP17 and FCHo1, and N-BAR domain from Endophilin-1. **b**. TIRF micrographs and kymographs of traveling waves of a representative RBL cell co-overexpressing GFP-FBP17 (**left)** and N-BAR (Endophilin-1^1-247^)-mCherry (**right)**. **c**. Intensity profile of GFP-FBP17 (**green**) and N-BAR (Endophilin-1^1-247^)-mCherry (**magenta**) of the same ∼2×2 µm^2^ region-of-interest (ROI) inside the cell in **b**. **d**. Cross-correlation (Xcorr throughout all figures) of the two signals in **c** shows the phase shift of waves of GFP-FBP17 relative to that of N-BAR (Endophilin-1^1-247^)-mCherry. **e**. Kymographs (**left**), intensity profile of the ∼2×2 µm^2^ ROI (**middle**), and cross-correlation (**right**) of waves of F-BAR(FCHo1^1-275^)-GFP (**green**) relative to that of mCherry-FBP17 (**magenta**). **f**. Intensity profile (**left**), and cross-correlation (**right**) of F-BAR(FBP17^1-329^)_K66E-GFP relative to mCherry-FBP17. Time resolution: 0.21s. **g**. Same as **f**, but with F-BAR(FBP17^1-329^)_K166A-GFP. All Scale bar: 10 µm. Time resolution: 1 s (**c, e**), 0.21 s (**f, g**),

While membrane curvature is likely continuously generated, sequential recruitments of curvature-preferring proteins can only happen if membrane curvature is the rate-limiting step for their recruitment and the rate of membrane bending is sufficiently slow to allow curvature-sensing. We further tested the curvature progression model by examining proteins showing even shallower curvature preferences than F-BAR^FBP17^. The F-BAR domain (a.a. 1-275) from FCHo1 (F- BAR^FCHo1^) induces membrane tubules with larger diameter than F-BAR^FBP17^ (Henne et al., 2007). We found that it was recruited to membrane waves 3 s before FBP17 (**Fig. 1e**). This relative timing is independent of the fluorescent tags used (Comparing between **Fig.1e** and **Supp Fig. 1d, e**) and imaging acquisition intervals (**Supp Fig. 1d, e**). To test whether the phases of these waves could reveal even smaller differences in curvature, we compared the effects of point mutations K66E and K166A in F-BAR^FBP17^. K66E mutant generates wider tubules *in vitro*, while K166A mutant generates narrower tubules than wild type F-BAR (Frost et al., 2008), but both to a lesser degree than N-BAR or F-BAR^FCHo1^. Using fast stream acquisition mode at 5Hz (0.2 s per frame), we found that F-BAR^K66E^ shifted the phase ahead of FBP17 by about 0.6 s (three frames of imaging) and F-BAR^K166A^ delayed the phase by about 0.4 s (**Fig. 1f**). Collectively, these phase shifts were consistent with our hypothesis that the curvature preference of proteins could predict their timing in recruitment to membrane waves.

### Two isoforms of FBP17 differ in their phases of recruitment to waves

FBP17 has two major isoforms. The short one is referred as FBP17 and the full-length is FBP17L throughout this paper (Kakimoto et al., 2004) (**Fig. 2a**). We co-expressed FBP17 and FBP17L in the same cell and found that both isoforms were recruited to the waves (**Fig. 2b**). Interestingly, we consistently found a phase difference between them, with FBP17 preceding FBP17L by 0.8-1.0 s, as quantified by cross-correlation analysis (**Fig. 2c**). We also confirmed that the relative phase differences between these two isoforms did not depend on which fluorescent tag combinations were used (**Fig. 2d, e**) or whether the fluorescent tags were at N- or C-terminus (**Supp Fig. 2a, b**).

**Figure 2.**
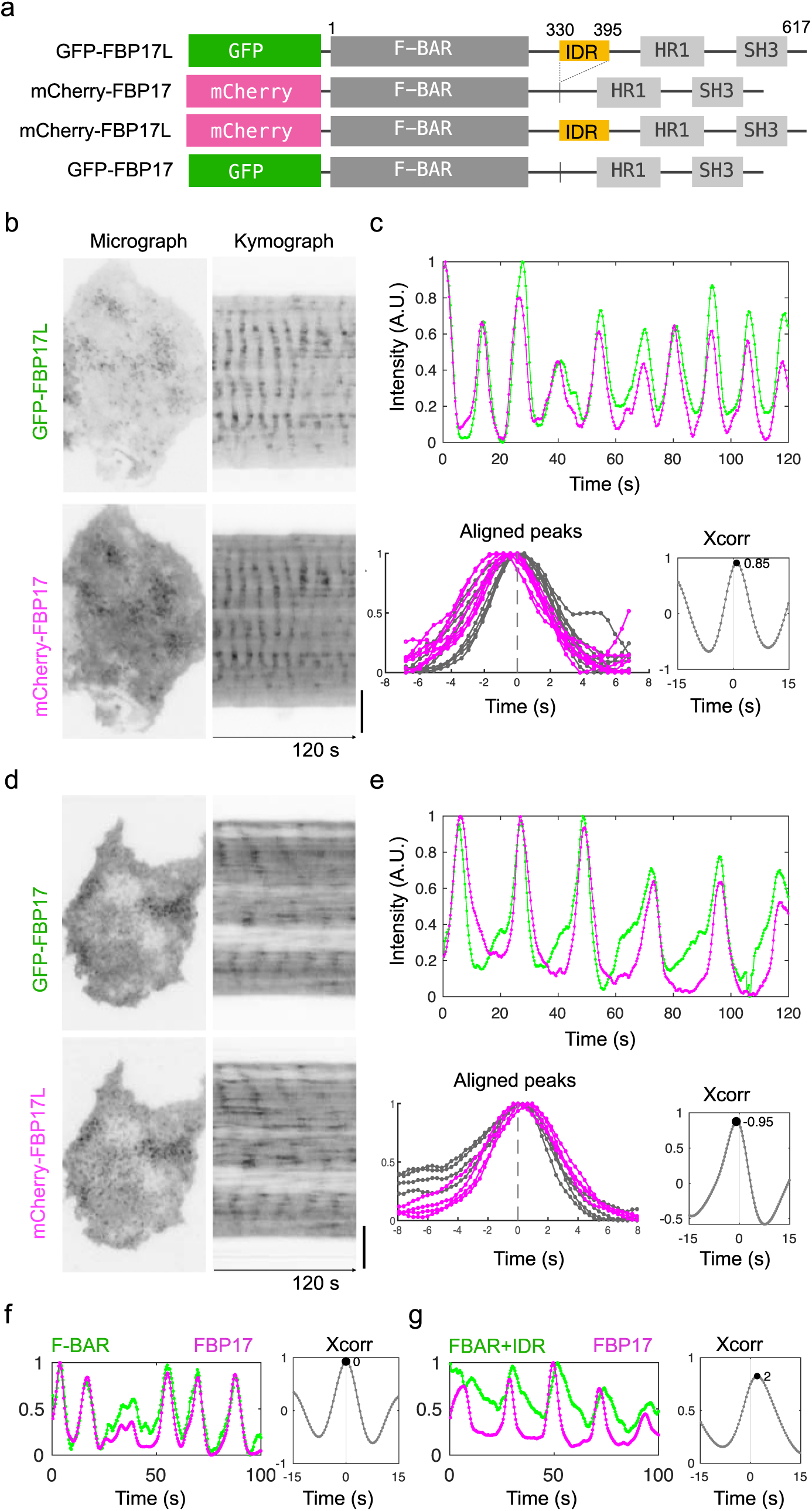
Phase shifts between FBP17 isoforms in traveling waves. **a**. Diagram of domain structures of full-length FBP17 (FBP17L) and short isoform (FBP17) with fluorescence tags fused at N-terminus. The intrinsic disordered region (IDR) is shown in yellow. **b,d**. Micrographs (**Left**) and kymographs (**right**) of co-overexpressed FBP17 in RBL cells. **c,e. Top**: Intensity profile of GFP (**green**) and mCherry (**magenta**) channels. **Bottom left**: aligned peaks of the green channel (shown in grey) from the **top** plots. Signal in mCherry channel (**magenta**) was plotted based on the same alignment in GFP channel. **Bottom right:** Cross-correlation of GFP relative to mCherry channel. **f,g**. Intensity profile (**left**), and cross-correlation (**right**) of F-BAR (FBP17^1-329^)-GFP (**f**) and F-BAR+IDR (FBP17^1-395^)-GFP (**g**) relative to FBP17-mCherry. Scale bar in **b,d**: 10 µm. Time resolution: 0.42 s (**b, c**), 0.32 s (**d, e**), 0.40 s (**f,g)**.

To test whether the observed phase differences were due to the membrane-binding difference between FBP17 and FBP17L, or more complex mechanisms such as conformational changes involving domains at C-terminus and other protein recruitment, we deleted the C-terminal a.a. 396- 617 (containing HR1 and SH3 domains) from both FBP17 and FBP17L. We co-expressed the remaining F-BAR domain alone (1-329 a.a. from FBP17) only in the same cells with FBP17. The F-BAR domain participated in the waves at the exact same phase of FBP17 (**Fig. 2f**), suggesting that the phase of FBP17 was set by the F-BAR domain, not its C-terminal domains. However, a.a. 1-395 from FBP17L (later named as F-BAR+IDR) (**Supp Fig. 2a**) was recruited about 2 s later than FBP17 (**Fig. 2g**). The phase lag was also independent of the time interval we chose, as long as it was less than half of the phase lag (**Supp Fig. 2c, d**). We thus concluded that the observed phase differences were due to the presence of the a.a. 330-395 following F-BAR domain in FBP17L.

### FBP17L contains conserved alternatively spliced intrinsic disordered region (IDR)

The shorter isoform FBP17 lacked a.a. 330-395 compared to FBP17L. This region in FBP17 is alternatively spliced (**Fig. 3a**). It is predicted to be intrinsically disordered and low in protein binding (**Fig. 3b**). We refer to the region a.a. 330-395 of human FBP17L as IDR^FBP17L^ in this paper. Interestingly, unlike many other IDRs (van der Lee et al., 2014), the amino acid sequence of the IDR^FBP17L^ is evolutionarily conserved. We found that the protein FBP17 appeared in the phylum Chordata and Arthropoda according to GeneBank and the exact sequence of the IDR of human FBP17 was found in the Clade Amniota comprising the reptiles, birds, and mammals (see Methods). To see the sequence conservation, we generated the sequence logo (Crooks et al., 2004) from all FBP17 sequences (**Fig. 3c**). At each position, the overall height (entropy) indicates the sequence conservation, while the width (weight) is proportional to the fraction of existence. IDR has few variations as shown by the alignment between the IDR of human FBP17 and consensus between IDRs of FBP17 from all species. For comparison, the paralog of FBP17, CIP4 (TRIP10), shares a sequence similar to FBP17 (71% positive for full-length and 66% for the corresponding IDR) (**Supp Fig. 3a**) but the sequence logo of its IDR (IDR^CIP4^) (**Fig. 3d**) is lower in entropy and weight (P<0.0001, Mann–Whitney U-test) (**Supp Fig. 3c**), indicating that the IDR^FBP17L^ is more evolutionarily conserved than IDR^CIP4^. The other paralog Toca-1 (FNBP1L) was also analyzed and its IDR is more conserved than IDR^CIP4^, but its weight is less than IDR^FBP17L^, indicating a higher likelihood to be spliced (**Supp Fig 3b, c**).

**Figure 3.**
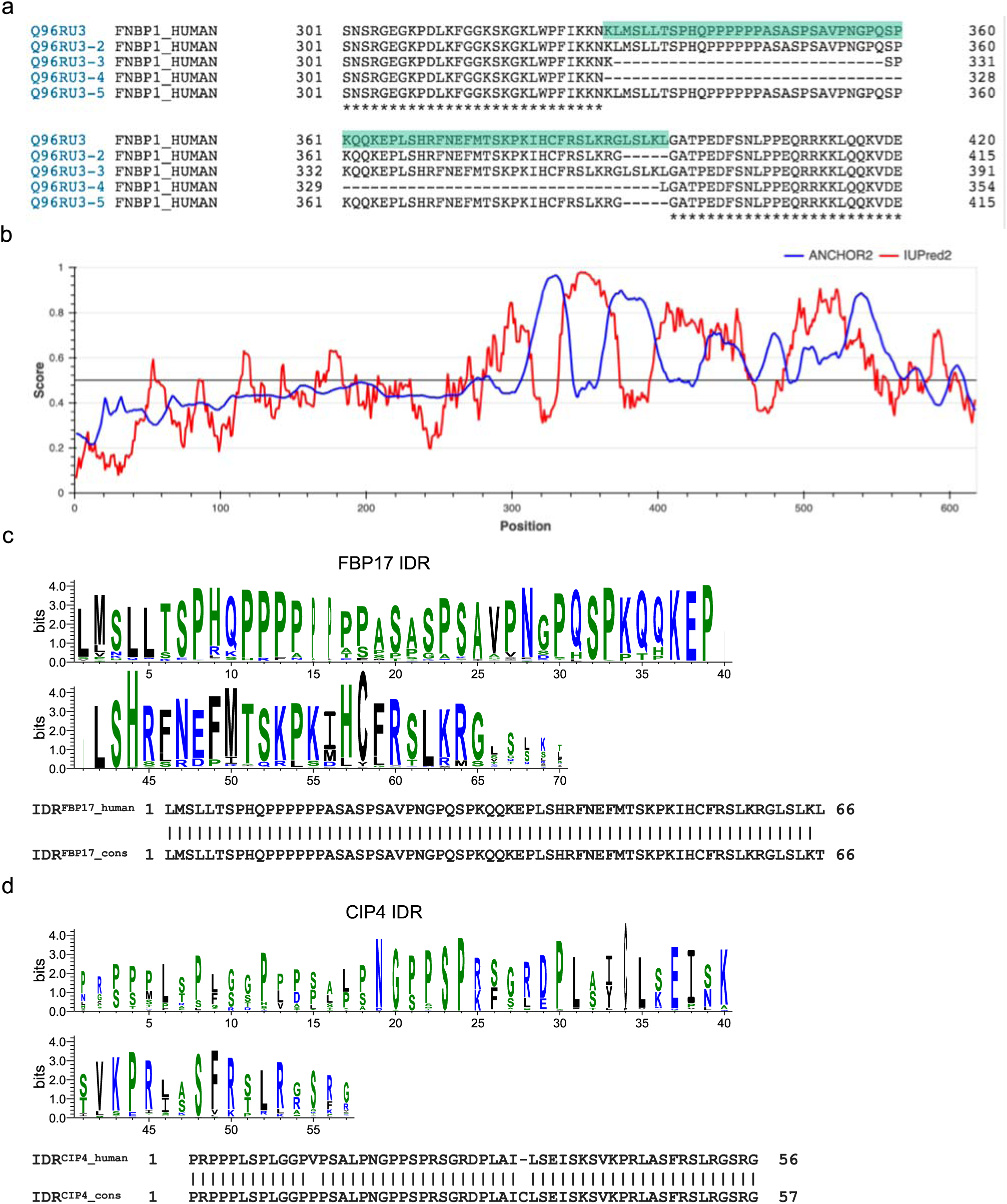
FBP17L contains a conserved intrinsically disordered region (IDR). **a**. Aligned alternatively spliced isoforms of human FBP17. The IDR is highlighted. **b**. Intrinsic disorder prediction of human FBP17L by IUPred2A, and protein-binding prediction by ANCHOR2. **c. Top:** WebLogo of the IDR of FBP17 from all 299 species that have FBP17. Blue: hydrophilic a.a. Green: neutral a.a. Black: hydrophobic a.a. **Bottom**: Sequence alignment of the IDR from human FBP17 and the consensus sequence from WebLogo3. **d. Top:** WebLogo of the IDR of CIP4 from all 183 species that have CIP4. **Bottom**: Sequence alignment of the IDR from human CIP4 and the consensus sequence from WebLogo3.

### IDR of FBP17L directly senses membrane curvature *in vitro*

While our results in living cells suggest a possible role of IDR in curvature-sensing, IDRs are flexible and subject to post-translational modifications in complex cellular context. It would be difficult to dissect a direct role of curvature sensing. We therefore set up an *in vitro* system based on the newly developed nanobar platform to directly test curvature sensing of the IDR and F-BAR from human FBP17L (Li et al., 2019; Zhao et al., 2017). As illustrated in **Fig. 4a**, one nanobar contained two curved ends with curvature defined by the bar width, while the flat middle served as local zero-curvature control. The scanning electron micrograph (SEM) (**Fig. 4a)** shows two arrays of vertically aligned nanobars of 300 nm in width, 2 µm in length and 600 nm in height fabricated via electron beam lithography. To study the differential binding of proteins on curved membrane, a supported lipid bilayer (10% brain phosphatidylserine, 0.5% Texas-Red DHPE in egg phosphatidylcholine) was formed on nanobars. The lipid bilayer covered the whole nanobar but did not show enrichment at the ends, indicating that the lipid coating was uniform (**Fig. 4c, left)**. We purified both IDR and F-BAR and tested their curvature preferences in the nanopillar- supported lipid bilayer system (**Fig. 4b**). The IDR showed stronger fluorescent signals at nanobar ends in comparison with nanobar centers (**Fig. 4c, right**). F-BAR also bound preferentially to curved nanobar ends, but to a lesser extent (**Fig. 4c, middle**). The difference in curvature preferences between IDR and F-BAR is more obvious in images averaged over 100 nanobars (**Fig. 4d**). Using the ratio of fluorescence intensities of nanobar end to center as an indicator of curvature preference (**Fig. 4a**), significant differences between IDR(1.4 ± 0.011) and F-BAR (ratio of 1.1 ± 0.004) were observed (unpaired t-test, p<0.0001), with either of them giving a higher ratio than lipid bilayer (1.0 ±0.001) (unpaired t-test, p<0.0001) (**Fig. 4d, e**). It shows that the IDR or F-BAR from human FBP17L alone is capable of sensing curved membrane directly.

**Figure 4.**
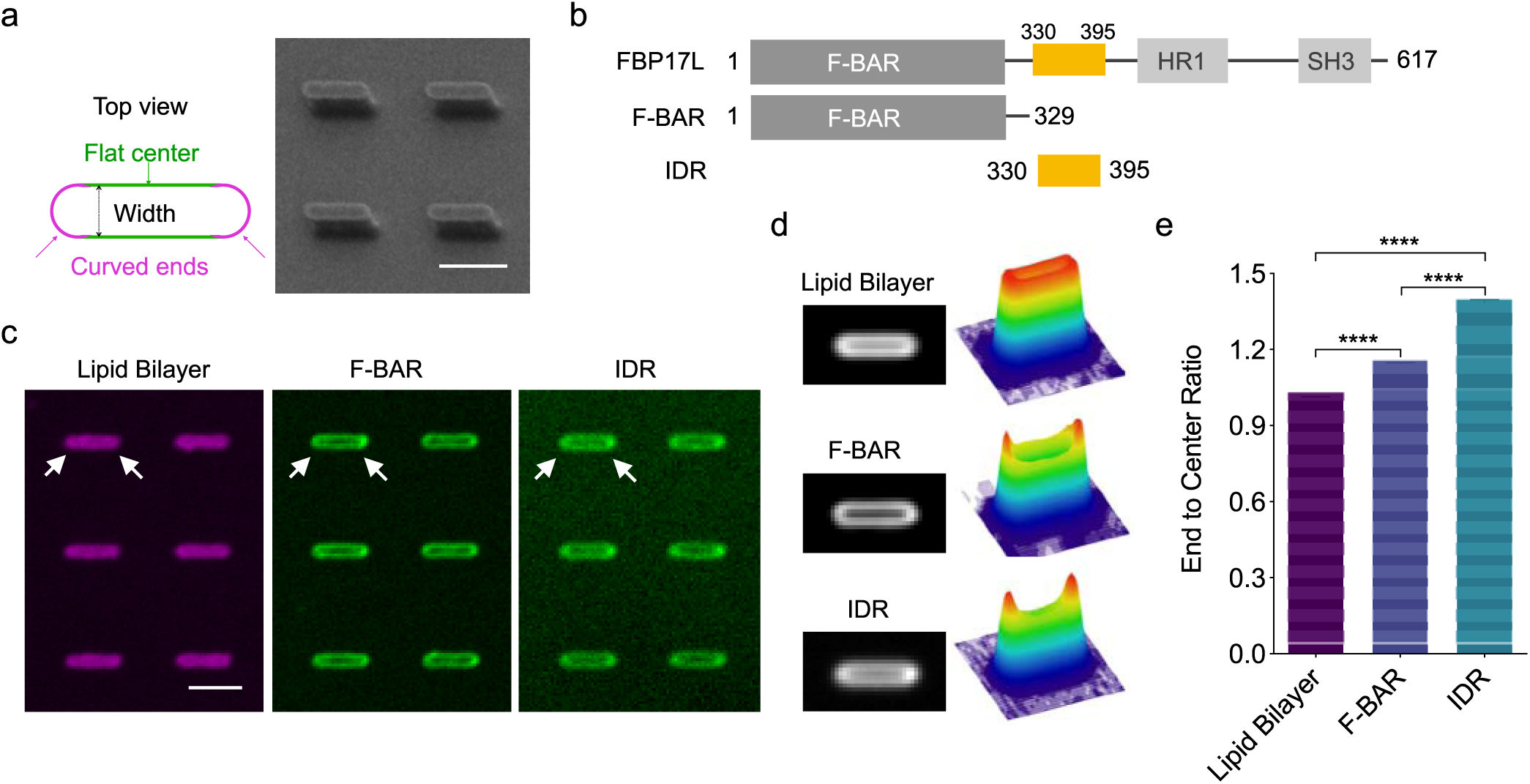
IDR of FBP17L senses membrane curvature *in vitro*. **a**. Schematic illustration of nanobar structure with combined curvature design and the scanning electron micrography (SEM) image of fabricated nanobar array. Scale bar: 2 µm. **b**. Diagram of domain structures of FBP17L and its truncations: F-BAR (a.a. 1-329) and IDR (a.a. 330-395). **c**. Confocal images of Texas Red labelled lipid bilayer, 16 µM Alexa Fluor 488 labelled F-BAR and 16 µM Alexa Fluor 647 labelled IDR coated on nanobars of 300 nm width. Arrowheads indicate two nanobar-ends. Scale bar: 2 µm. **d**. Average images and the corresponding 3D surface plot of lipid bilayer, F-BAR and IDR on over 100 nanobars of 300 nm width. **e**. End to center ratio of lipid bilayer, F-BAR, and IDR on 200 nanobars of 300 nm width (n=3 experiments per protein). Unpaired t-test with welch’s correction: **** p<0.0001. Graph shows mean ± s.e.m.

To examine the range of membrane curvatures that the IDR senses, we employed the nanobar array with a series of defined sizes. The nanobars with bar-end curvature diameters varying between 200 nm to 600 nm were patterned in arrays in 100 nm increments (**Fig. 5a**). Compared to the homogeneous lipid coating on all nanobars with different widths (**Fig. 5b**), the IDR showed higher signals at the nanobar ends as the diameter of the bar-ends decreased to below 400nm (**Fig. 5c**). In comparison, the corresponding IDR^CIP4^ had no detectable binding to the lipid bilayer at any of the sizes tested (**Fig. 5d**), suggesting a unique role of IDR^FBP17L^ in curvature sensing. Similar to the IDR, F-BAR and the combined F-BAR+IDR also showed increased signals at bar-ends (**Fig. 5e, f**). To compare the relative degrees of enrichment across nanobars of different sizes, we normalized the bar-end intensity for each protein with that of the fluorescent lipids to account for any geometric effects. All three proteins showed stronger binding preference at higher membrane curvatures (**Fig. 5g**), with IDR showing the highest curvature response, F-BAR the lowest and F-BAR+IDR at an intermediate level.

**Figure 5.**
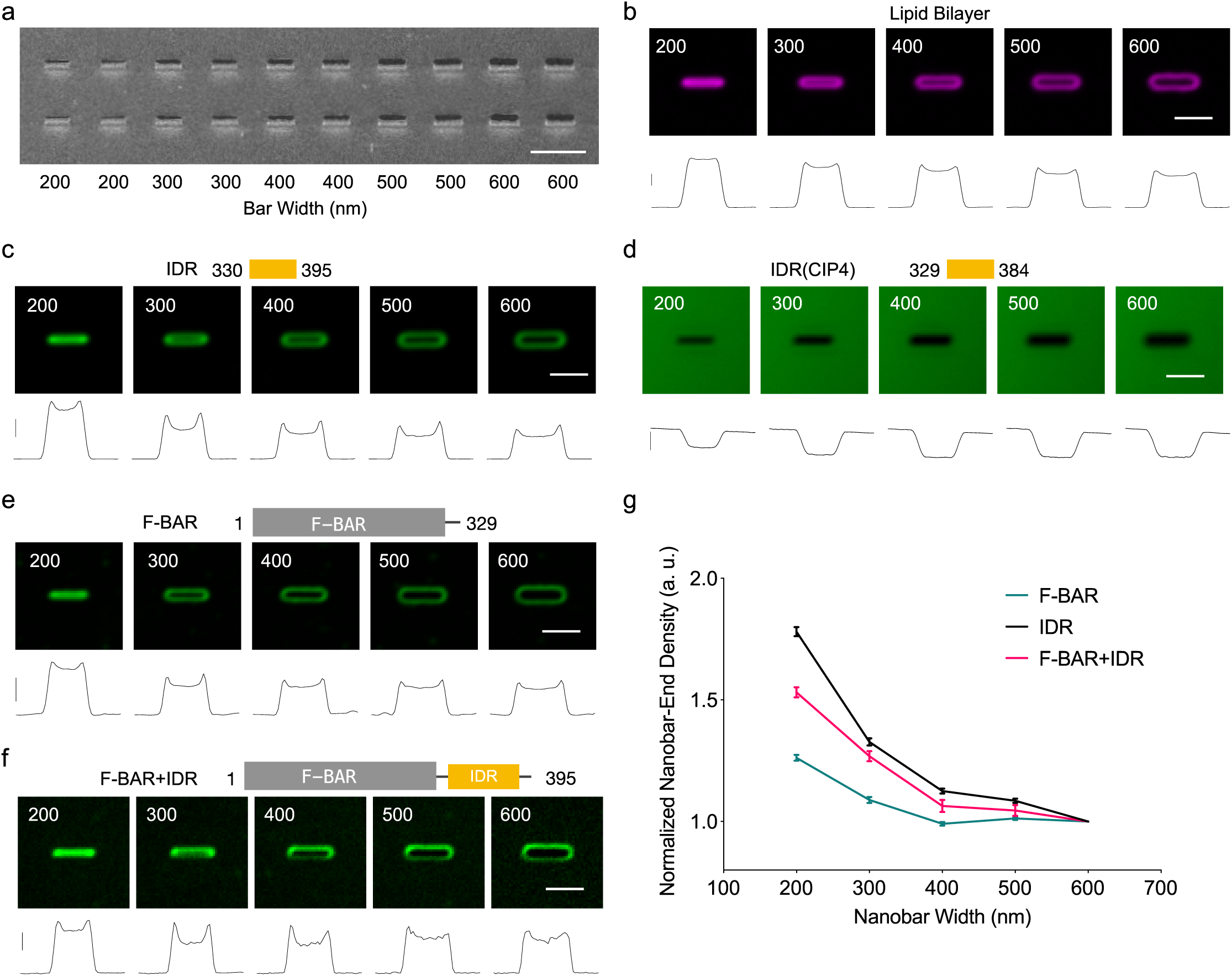
Curvature sensing of IDR, F-BAR, F-BAR+IDR on nano bar array. **a**. SEM image of nanobar arrays in width gradient from 200 nm to 600 nm with 100 nm step. Every two columns of the arrays are the same size as indicated. The height of every nanobar is 600 nm. Scale bar: 5 µm. **b-f**. Diagrams (**top**), averaged images (**middle**) of lipid bilayer (**b**), 16 µM IDR (**c**), 25 µM IDR(CIP4) (**d**), 16 µM F-BAR (**e**) and 16 µM F-BAR+IDR (**f**) on nanobars with width from 200 nm to 600 nm. Each image was averaged from over 60 nanobars. Scale bar: 2 µm. **Bottom**: Line profiles of each averaged image. The x-axis represents horizontal distance through nanobar and the y-axis is the vertically averaged top 25% intensity within nanobar area. Vertical scale bars at the left: from **b** to **f**: 5000 a.u., 2000 a.u., 1000 a.u., 5000 a.u., 5000 a.u.. **g**. Normalized nanobar end density of F-BAR, IDR and F-BAR+IDR based on their corresponding lipid bilayers intensity. Each point represents mean from over 60 nanobars and the error bar represents s.e.m (n=3 experiments per protein).

### Faster dissociation rate of the IDR compared to F-BAR domain *in vitro*

We next compared the dynamics of membrane binding of IDR or F-BAR using fluorescence recovery after photobleaching (FRAP). After bleaching a single nanobar area within a circle area of 5 µm diameter, the lipid bilayer quickly recovered with a half-time (t_1/2_) of about 9s (**Fig. 6a, e**), indicating good membrane lipid fluidity. For F-BAR, there is no detectable recovery within the same 2-min window of the experiment (**Fig. 6b, e**), consistent with the previously reported higher-order assembly of F-BAR domain formed via oligomerization on membrane (Frost et al., 2008). Interestingly, the IDR showed significant recovery with a half-time of about 35s (**Fig. 6c, e**). To further test whether the IDR recovery was due to membrane binding and dissociation or lateral diffusion, we washed away the unbound IDR with buffers and performed the same FRAP experiment. We observed no obvious recovery at the same time scale, suggesting that the lateral diffusion via curvature-based protein sorting contributed little to the recovery (**Fig. 6d, e**).

**Figure 6.**
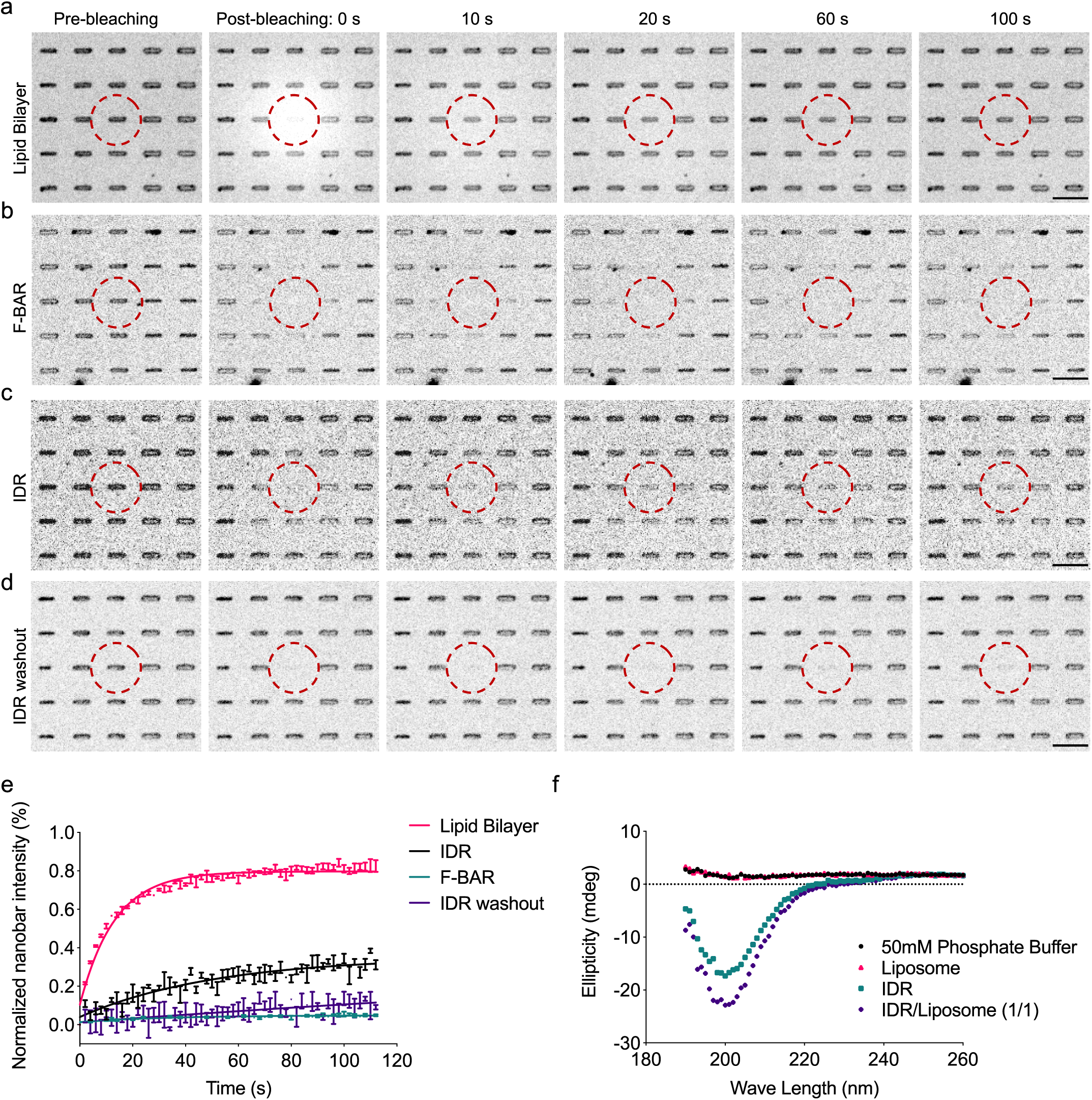
Membrane binding dynamics of IDR and F-BAR. **a-d**. Montages of confocal FRAP test on lipid bilayer (**a**), F-BAR (**b**), IDR (**c**) and IDR washed out (**d**) on nanobar with 300 nm width in 2 minutes. Red dashed circles indicated the bleaching area. Scale bar: 5 µm. **e**. Normalized intensity plot of lipid bilayer, F-BAR, IDR and IDR washed out on single nanobar from FRAP measurement (2 s interval) (n = 3 experiments per condition). The measurement showed lipid bilayer’s recovery with turnover rate at 0.076 ± 0.003 sec^-1^ and immobile fraction at 20 ± 0.4%. IDR recovered with turnover rate at 0.02 ± 0.004 sec^-1^ and immobile fraction at 65 ± 2%. **f**. Representative circular dichroism spectra of phosphate buffer, liposomes only, 14 µM IDR and IDR 1:1 mole ratio incubated with liposomes of ≤ 350 nm diameter at room temperature.

Previously, it was shown that disordered amphipathic lipid packing sensor (ALPS) can sense curvature by forming stable helix upon binding of the lipid bilayer (Drin et al., 2007). However, for the IDR, no folded structure was detected by circular dichroism (CD) measurement even after binding to liposomes (**Fig. 6f**), excluding the impact of potential folding and helix formation of IDR upon membrane binding. Nevertheless, we cannot rule out the possibility of hydrophobic interaction at the level of single amino acid at this stage.

### High concentration promotes curvature sensing of the IDR

Curvature sensing has been shown to either increase (Ramamurthi et al., 2009) or decrease (Ramesh et al., 2013; Zeno et al., 2018) with higher concentration, indicating different dependency of curvature sensing on protein proximity. To further dissect the IDR interaction with curved membranes, we compared the nanobar binding of the IDR at a series of concentrations. At higher concentration, more IDR bound to the nanobar, but this effect was stronger on smaller bars (**Fig. 7a)**. When the relative enrichment of IDR at nanobar ends was quantified, higher concentration clearly led to increased degree of curvature sensing (**Fig. 7b)**. Consistent with curvature sensing enhanced by protein proximity, binding curve of the IDR on membranes show stronger cooperativity compared to that of F-BAR or F-BAR+IDR (**Fig. 7c-e**). We could not reach saturating concentration for F-BAR or F-BAR+IDR, but for the IDR, we fitted its binding curves with the Hill equation, and obtained the Hill coefficient (*H*) between 2-3 and *K*_*D*_ (0.6 ± 0.01 μM), suggesting an ultrasensitive binding of IDR to highly curved membrane sites.

**Figure 7.**
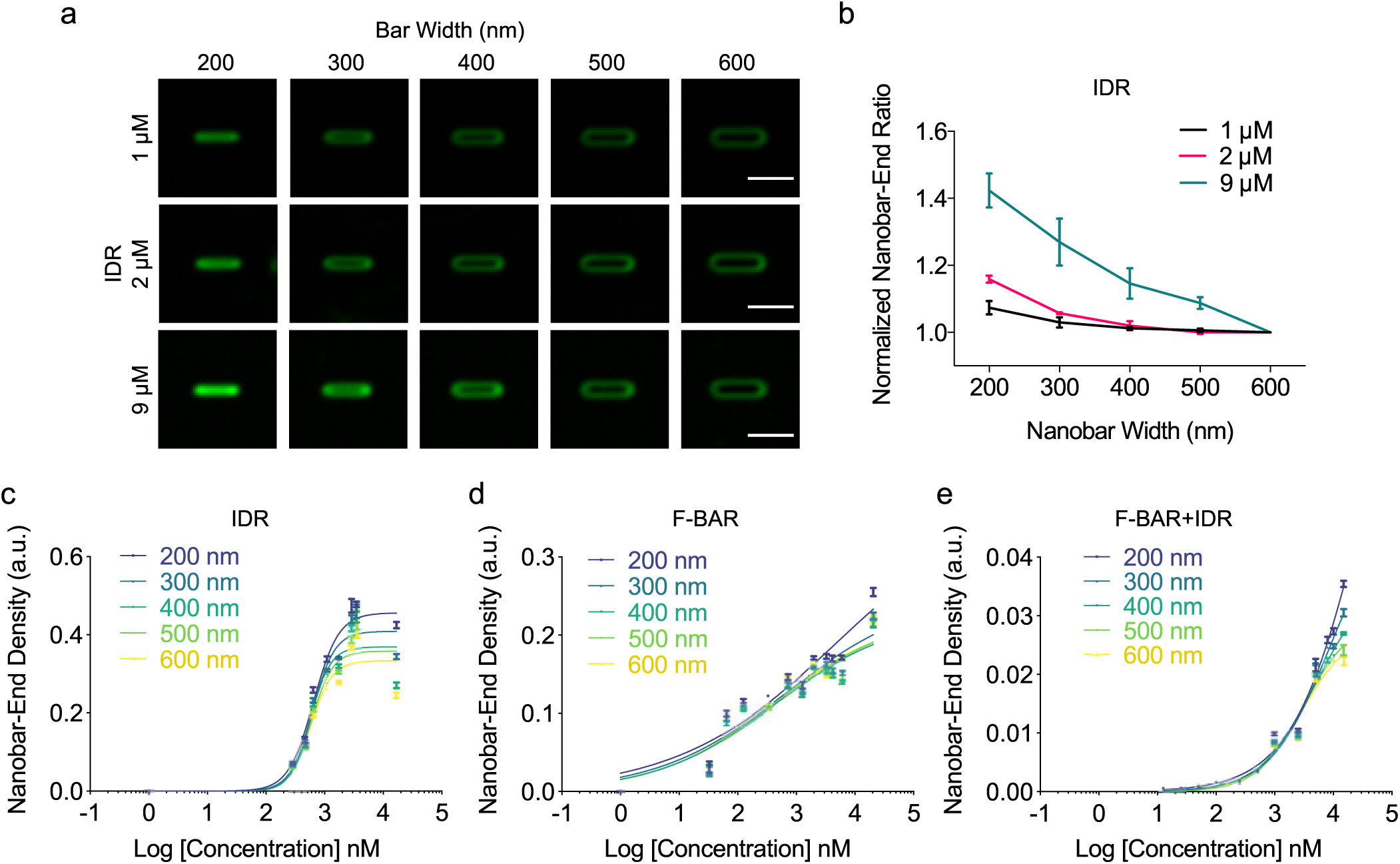
High concentration enhanced IDR’s curvature sensitivity. **a**. Averaged images of IDR at different concentrations on nanobars with gradient width from 200 nm to 600 nm. Each nanobar image was averaged from over 60 nanobars. Scale bar: 2 µm. **b**. Normalized nanobar end density of IDR at different concentration. Each point represents mean from over 60 nanobars and error bar represents s.e.m. **c-e**. Plot of nanobar-end density of IDR (**c**), F-BAR (**d**) and F-BAR+IDR (**e**) as a function of concentration. Lines are binding curves fitted with the Hill equation. Error bar represents s.e.m.

## Discussion

In this work, by carefully examining the phases of the protein waves, we discovered a conserved IDR in FBP17L that affected the kinetics of protein recruitment to sites of membrane bending in living cells. This motivated us to examine *in vitro* whether IDR could directly sense curvature. Certainly, a direct role in curvature sensing could only be demonstrated through *in vitro* studies because lipid-binding modules as well as IDRs are likely subjected to additional regulations in cells. Indeed, we found that the curvature sensitivity of the F-BAR domain is modulated by this adjacent IDR. Current understanding of the mechanism of membrane curvature sensing is predominantly through well-defined domain structures that have membrane insertion (e.g. ENTH domain, amphipathic helices), curved membrane-binding surfaces (e.g. BAR domain) or both (e.g. N-BAR domain). F-BAR domains were widely assumed to sense curvature due to its crescent-shape, but the curvature sensitivity of F-BAR from FBP17 has never been demonstrated in the literature. The curvature sensing provided by IDR has several precedents. IDRs were proposed to play an important role in endocytic progression (Ungewickell and Hinrichsen, 2007). IDRs from endocytic proteins such as epsin, AP180 or amphiphysin were shown to generate curvature (Busch et al., 2015; Snead et al., 2019), as well as sensing curvature through molecular crowding (Zeno et al., 2018, 2019). Recently, the IDR of F-BAR protein Fer was also shown to sense curvature using liposome binding assay (Yamamoto et al., 2018). The IDR reported here is much smaller in size (66 amino acids) and does not require additional lipid anchors to recruit it to the membrane under our *in vitro* experimental conditions. We also provided a side-by-side comparison of the F-BAR domain and IDR from the same protein using a quantitative nanobar platform. At the same concentration, we observed stronger binding for the F-BAR domain alone but curvature sensing as measured by relative enrichment of end binding is stronger with the IDR. Given that the IDR could dynamically associate with membrane and mediate curvature sensing, this strongly indicates curvature sensing mediated by the IDR as a collective effect not dependent on imposing a rigid molecular scaffold over a curved surface.

We found the sequence of recruitment of F-BAR proteins FBP17, CIP4, N-BAR proteins Endophilin and Amphiphysin in the cortical waves depends on their membrane binding domains. Together with *in vitro* results confirming curvature sensing, we established that the sequential recruitment of these proteins in the travelling waves of mast cell cortex correlated with the progression of membrane curvature. A model of sequential curvature generation was hypothesized for endocytosis when the crystal structure of F-BAR was first solved (Shimada et al., 2007). It was thought that the F-BAR domain could sculpt a flat membrane into shallow curvature, followed by BAR protein, leading to the formation of clathrin-coated buds. While it is an intuitive model, this is conflicted with observations that F-BAR proteins of the FBP17/CIP4/Toca subfamily are recruited at late stages of clathrin-mediated endocytosis (Taylor et al., 2011), and its contribution to endocytosis are likely morphologically independent from the maturation of clathrin-coated pits (Wu et al., 2010; Yang et al., 2017). Here we confirm that the sequential curvature generation model can be true but in a modified context from the original proposal. The dynamics of curvature generation covers a time period of about 8 seconds (F-BAR^FCHo1^ at -3 sec, FBP17 or F-BAR^FBP17^ at 0 sec, endophilin, N-BAR^endo^ or N-BAR^amph^ at 5 sec) which is much shorter than the time it takes to assemble an endocytic pit (30 sec to 1 min), and it corresponds to the late stage of the endocytosis. The rapid time scale is consistent with our previous characterization of membrane undulation using Surface Reflective Interference Contrast (SRIC) microscopy. As membrane waves propagate and oscillate with a period of 20-30 s, the rise phase of SRIC lasts around 10 s (Wu et al., 2018). While the phase differences are on the second time scale, it represents a significant fraction of the total time for membrane bending occurring in a living cell. Because of the time scale of this process, changes in bulk cytoplasmic protein concentrations can be minimal. Under these experimental conditions, we show that curvature preferences as defined by the end result of curvature-generation *in vitro* can be reflected as the kinetics of their recruitments, suggesting that curvature-dependent membrane waves represent a unique context to infer curvature-sensing.

### Limitations of the Study

It remains to be determined whether fine tuning of the curvature sensing and recruitment kinetics of F-BAR domain-containing proteins through the IDR will result in cellular level functional consequences. Because the IDR of FBP17L is precisely what defines the divergence of naturally occurring splicing isoforms, and the its sequence is highly conserved across species, we speculate that the IDR’s curvature sensing has functional implications. Interestingly, a recent work showed FBP17 isoform-specific function in neuronal morphology as well as an IDR-dependent membrane tubulation activity in cells. The IDR domain of FBP17 was found to be more potent in inducing neurite formation (Taylor et al., 2019). Consistent with our findings, such property of IDR of FBP17 is not shared by the IDR in CIP4. In addition, the IDR of the cytokinetic F-BAR protein Cdc15 in fission yeast was shown to be important for the maintenance of cytokinetic ring (Mangione et al., 2019). Cell protrusions and leading edge dynamics as well as cytokinesis are often simplified as actin cytoskeletal-centric rearrangement but there are also controlled by cortical propagating waves and interlinked oscillatory networks that remain to be elucidated (Bement et al., 2015; Bolado-Carrancio et al., 2020; Katsuno et al., 2015; Miao et al., 2017; Ruthel and Banker, 1999; Xiao et al., 2017). An intriguing possibility is that these kinetic parameter changes could have implications in the conversion between regimes of dynamical systems (also known as bifurcations) which then result in morphogenetic outcomes. Information obtained here therefore would be expected to contribute to physical insights towards building a more holistic model of cellular dynamics.

## Methods

### Materials

L-α-phosphatidylcholine (Egg PC, Chicken, 840051) and L-α-phosphatidylserine (Brain PS, Porcine, 840032) were from Avanti Polar Lipids, Inc. Texas Red® DHPE was from Invitrogen™. IDRs of FBP17L and CIP4 (89% purity) were synthesized by Sangon Biotech (Shanghai) Co., Ltd. and GL Biochem (Shanghai) Ltd. The molecular weight was confirmed by HPLC and mass spectrometry. F-BAR^1-329^ and F-BAR+IDR^1-395^ of FBP17L were expressed and purified as fusion proteins with His-Flag tag in *E. coli* BL21 (DE3) Rosetta T1R by NTU Protein Production Platform (www.proteins.sg).

### Plasmids

mCherry-FBP17L, FBP17-EGFP and chimeric Endo1-FBP17-GFP were described previously (Wu et al., 2018). EGFP-FBP17L and FBP17L-mCherry were acquired by cloning human FBP17 (UniProt ID: Q96RU3, full length, a.a. 1-617) into pEGFP-C2 and pmCherry-N1 vectors. FBP17L-EGFP was generated by adding a.a. 330-390 back through PCR-ligation to FBP17-EGFP. EGFP-FBP17 and mCherry-FBP17 were generated by deleting a.a. 330-395 from EGFP-FBP17L and mCherry-FBP17L. FBP17-mCherry was obtained by cloning FBP17 from FBP17-EGFP to pmCherry-N vector. F-BAR from FBP17 (a.a. 1-329), IDR of FBP17L (a.a. 330-395), F-BAR+IDR from FBP17L (a.a. 1-395), F-BAR from human FCHo1 (a.a. 1-275), N-BAR from human Endophilin A1 (a.a. 1-247), N-BAR from human Amphiphysin1 (a.a. 1-236) were cloned to pEGFP-N1 or pmCherry-N1 to generate respective domain-only plasmids. Endophilin-2-GFP was a gift from Pietro De Camilli. Point mutations (K66E and K166A) on F-BAR-EGFP were produced by the Q5 Site-Directed Mutagenesis Kit (New England BioLabs). Identity of plasmids was confirmed by Sanger sequencing.

### Cell Culture and Transfection

As described previously (Wu et al., 2018), RBL-2H3 cells (tumor mast cells) were maintained in MEM medium with 20% FBS. Cells were harvested with TrypLE (Life Technologies) 2–5 days after passage. Electroporation with Neon transfection system (Life Technologies) was utilized for transient transfections. After transfection, cells were plated at subconfluent densities in 35-mm glass bottom dishes (Cellvis, CA) overnight. Cells were sensitized with mouse monoclonal anti- 2,4-dinitrophenyl IgE (Sigma-Aldrich) at 0.5 μg ml^−1^ overnight and stimulated with 80 ng ml^−1^ multivalent antigen, 2,4-Dinitrophenylated bovine serum albumin (Life Technologies) before imaging. During imaging, the cells were either in medium or Tyrodes buffer (135 mM NaCl, 5.0 mM KCl, 1.8 mM CaCl2, 1.0 mM MgCl2, 5.6 mM glucose, and 20 mM Hepes (pH 7.4)).

### TIRF Imaging and Image Analysis

The TIRF microscopy imaging system was described previously (Wu et al., 2018). In brief, a Nikon Ti-E inverted microscope, iLAS2 motorized TIRF illuminator (Roper Scientific, Evry Cedex, France) and Prime 95B CMOS Camera (Teledyne Photometrics, AZ) was used. All images were acquired through a TIRF-CFI objective (Apochromat TIRF 100XH numerical aperture (NA) 1.49, Nikon). For stream acquisition mode, the exposure time was set at 50 ms per channel to achieve 0.21 s frame interval (instead of 100 ms at 0.7 or 1.0 s interval). Images were analyzed by Fiji-ImageJ to generate micrographs, kymographs, and movies. Plots and graphs were generated in MATLAB 2018b. Phase shift was calculated by custom code of cross-correlation in MATLAB. A ∼2×2 μm^2^ region-of-interest (ROI) was selected at an area where waves were prominent and last for at least three cycles. The corresponding intensity profile vs time was smoothed (span=5) and normalized to 0-1 range. The phase shift was extracted at the maximum cross-correlation while shifting signal from channel 1 (EGFP) relative to channel 2 (mCherry) at -15 s to +15 s range. The phase shift at maximum correlation > 0.6 was shown. Statistics and plots were performed by the built-in functions of PRISM 8.

### Bioinformatics

The sequences of all FBP17 and all CIP4 were extracted from UniProtKB by the search term ‘gene_exact:fnbp1’ and ‘gene_exact:trip10’. Sequence alignment was performed by Clustal Omega and the sequence logo with its entropy and weight data was generated by WebLogo 3.6.0. Intrinsic disorder prediction was performed by IUPred2A (Mészáros et al., 2018). The distribution of the IDR in the tree of life was produced by lineage and taxonomy report of Blastp (protein-protein BLAST) result using the human IDR sequence against database Non-redundant protein sequences. Each hit with a species had E value <10^−36^.

### Fabrication of Nanobar Chips

The nanobar arrays used in this study were fabricated on a square quartz wafer by using electron-beam lithography as described in previous work (Zhao et al., 2017). In brief, 15*15*0.2 mm square quartz coverslips were spin-coated with positive electron-beam resist PMMA (MicroChem) with around 300 nm height. Conductive protective coating was applied later with AR-PC 5090.02 (Allresist). Nanobar patterns were drawn by EBL (FEI Helios NanoLab) and developed in IPA: MIBK=3:1 solution. A 100 nm-height chromium mask was generated via thermal evaporation (UNIVEX 250 Benchtop) and lifted off in acetone. Nanobars were subsequently synthesized by reactive ion etching with the mixture of CF_4_ and CHF_3_ (Oxford Plasmalab80). Scanning Electron Microscopy (FEI Helios NanoLab) imaging was performed after 10 nm chromium coating. Before usage, the nanobar chips were immersed in Chromium Etchant (Sigma-Aldrich) until the chromium masks were removed.

### Protein Labelling

Alexa Fluor™ 488 TFP ester dyes (A37563, Invitrogen™) were dissolved in DMSO at a concentration of 10 mg/ml and stored at -80 °C. F-BAR and F-BAR+IDR were diluted to 2 mg/ml in PBS. 1 M sodium bicarbonate solution was prepared and added to the protein solution at a final concentration of 0.1M (pH∼8.3). 10 μl TFP ester dyes were warmed up to room temperature and transferred to the protein solution. The reaction mixture was inverted for 1 h at room temperature, resulting in a labeling ratio between 1:1 to 1.5:1 (dye:protein). Unconjugated TFP ester dyes were separated from labeled proteins using HiTrap Desalting columns with Sephadex G-25 resin (GE Healthcare Life Sciences). Alexa Fluor™ 647 C2 Maleimide (A20347, Invitrogen™) was used to label the only cysteine of IDR^FBP17^. For IDR^CIP4^, additional cysteine was added at the C-terminus during synthesis. Ligation was at room temperature for 30 min and 100 mM glutathione was added to stop the reaction. Unconjugated Maleimide dyes were dialyzed from labeled IDR using 3.5 kDa membrane in 1 L PBS overnight twice at 4°C. Amicon Ultra-0.5 mL Centrifugal Filters (Merck) were used to concentrate the labeled proteins. The final protein concentration and labeling ratio were measured using NanoDrop™ UV-Vis Spectrophotometer (Thermo Scientific™).

### Preparation of Lipid Vesicles

The molar composition of lipid vesicles is egg PC (Avanti) mixed with 0.5 mol% of Texas Red- DHPE (Invitrogen™) and 10 mol % of brain PS (Avanti). Lipids were dissolved in chloroform and mixed at a fixed molar ratio as described above. The lipid mixture was dried down in a brown glass vial with 99.9% nitrogen gas for 5 min, followed by vacuum drying for 3 h to remove the remaining chloroform. The dried lipid film was resuspended in PBS at a concentration of 2 mg/ml and sonicated for 30 min. The lipid mixture was then transferred to a 1.5 ml tube and freeze-thawed for 15 times (20 s in liquid nitrogen and 2 min in 42°C water bath). The lipid mixture was extruded through a polycarbonate membrane with 100 nm pore size using an extruder set with holder/heating block (610000-1EA, Sigma-Aldrich). The lipid vesicle solution was stored at 4 °C and used within 2 weeks.

### Lipid Bilayer Formation on Nanobar Chips

The nanobar chips were cleaned with piranha solution (7 parts of concentrated sulfuric acid and 1 part of 30% hydrogen peroxide solution) overnight and washed with a continuous stream of deionized water to remove the acid molecules. Subsequently, the nanobar chips were dried in 99.9% nitrogen gas and cleaned with air plasma in a plasma cleaner (Harrick Plasma) for 30 min to remove any remaining impurities on the surfaces before lipid bilayer formation. The cleaned chips were attached with a PDMS chamber and lipid vesicles were loaded into the PDMS channel and incubated for 15 min to form the lipid bilayer. PBS was finally used to wash away the unbound vesicles in the PDMS channel. Protein solution was subsequently added onto the lipid-bilayer-coated nanobar and incubated for 5 min before confocal imaging at room temperature.

### Imaging of Protein-Nanobar Interaction and Quantification

Imaging of purified proteins distribution on lipid-bilayer-based nanobar structures was performed with laser scanning confocal microscopy (Zeiss LSM 800 with Airyscan) using 100x/1.4 oil objective. Excitation of Alexa Fluor™ 488-labeled F-BAR and F-BAR+IDR was performed at 488 nm and detected at 519 nm. Excitation of Alexa Fluor™ 647-labeled IDR was performed at 633 nm and detected at 665 nm. Each image had a resolution of 512×512 pixels, with a pixel size of 124 nm and 16-bit depth. The intensity of nanobars was measured and calculated using a custom-written MATLAB code as previously reported (Zhao et al., 2017). Statistics and plots were performed by PRISM 8.

## Acknowledgements

We thank Cheesan Tong, Su Guo, Jeffery Yong, Jiaxi Cai, and Jianwei Li (Adam Yuan lab) for experimental assistance, Jianxin Song, Yansong Miao, Jian Liu for discussion. We thank the NTU Protein Production Platform (www.proteins.sg) for protein purification. Fabrication is supported by the Nanyang NanoFabrication Center (N2FC) and the Centre of Disruptive Photonic Technologies (CDPT) in Nanyang Technological University. This work is supported by the National Research Foundation Singapore (NRF) under its NRF Fellowship Program (M. Wu, NRF award No. NRF-NRFF2011-09), the Singapore Ministry of Education (MOE) Academic Research Fund Tier 1 (W. ZHAO, 2018-T1-002-098) and Tier 2 (M. Wu, 2015- T2-1-122), the Singapore Ministry of Health National Medical Research Council Open Fund Individual Research Grant (M. Wu, NMRC/OFIRG/0038/2017), Yale University startup grant (M. Wu), Nanyang Technological University Start-up Grant (W. Zhao) and NTU-NNI Neurotechnology Fellowship (W. Zhao).

---

**Supplementary Figure 1.**
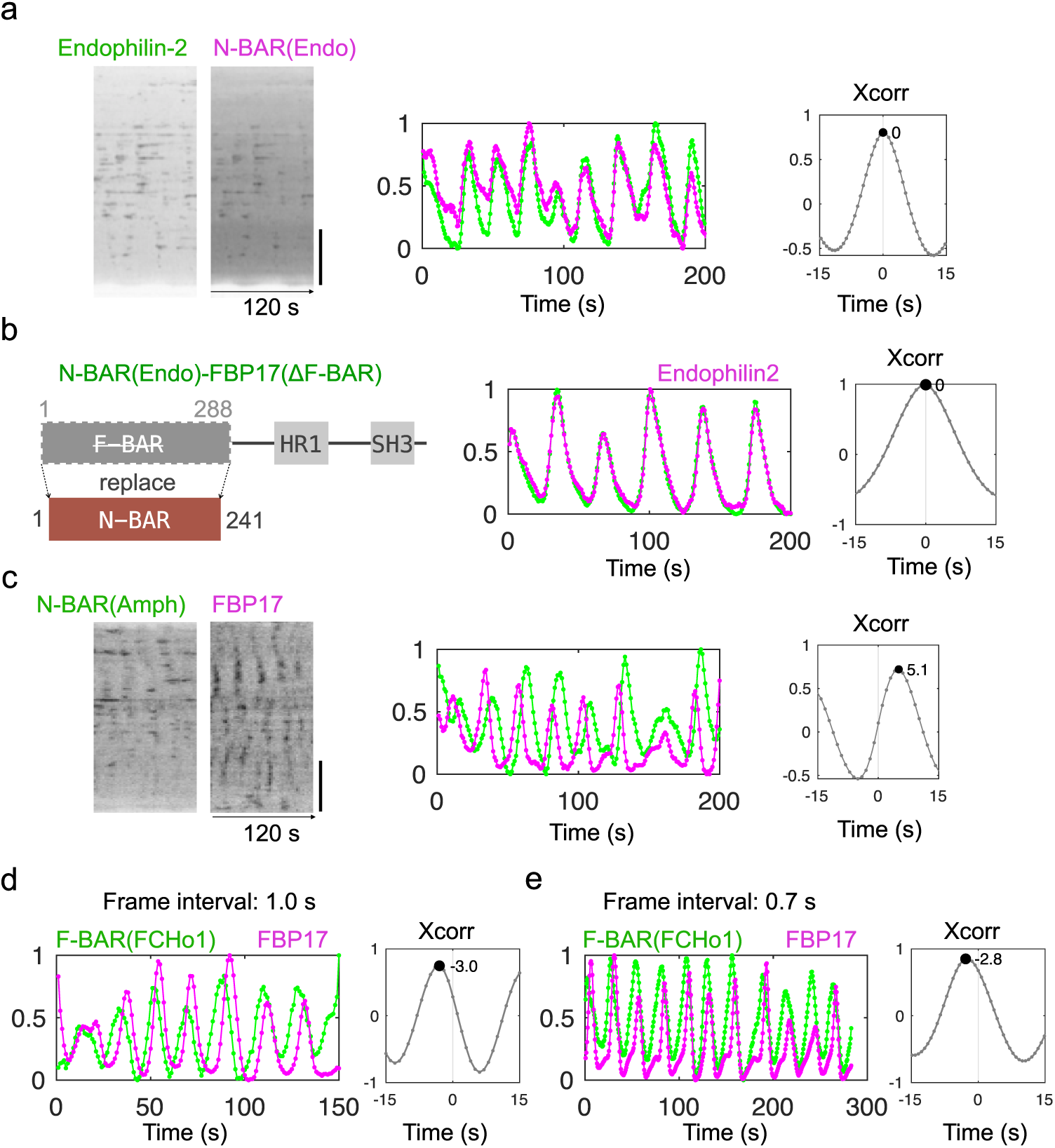
Timing difference in waves is integral to the BAR domains. **a**. Kymographs of the same line within cell area (**left**), intensity profile of the same ∼2×2 µm^2^ ROI in the cell (**middle**), and cross-correlation (**right**) of waves of Endophilin-2-GFP and N-BAR (Endophilin-1^1-247^)-mCherry. Time resolution: 0.7 s. Scale bar: 10 µm. **b. Left:** Diagram of domain structure of chimeric N-BAR(Endophilin-2^1-241^)-FBP17^ΔF-BAR^-GFP (not drawn to scale). Intensity profile (**middle**), and cross-correlation (**right**) of waves of it relative to waves of Endophilin-2-mCherry. Time resolution: 1 s. **c**. Kymographs (**left**), intensity profile (**middle**) and cross-correlation (**right**) of waves of N-BAR(Amphiphysin-1^1-236^)-GFP and FBP17-mCherry. Time resolution: 1 s. Scale bar: 10 µm. **d,e**. Phase shift of F-BAR (FCHo1^1-275^)-GFP relative to mCherry-FBP17 imaged at time interval of 1.0 s (**d**) and 0.7 s (**e**). **Left**: Intensity profile. **Right:** Cross-correlation of GFP relative to mCherry.

**Supplementary Figure 2.**
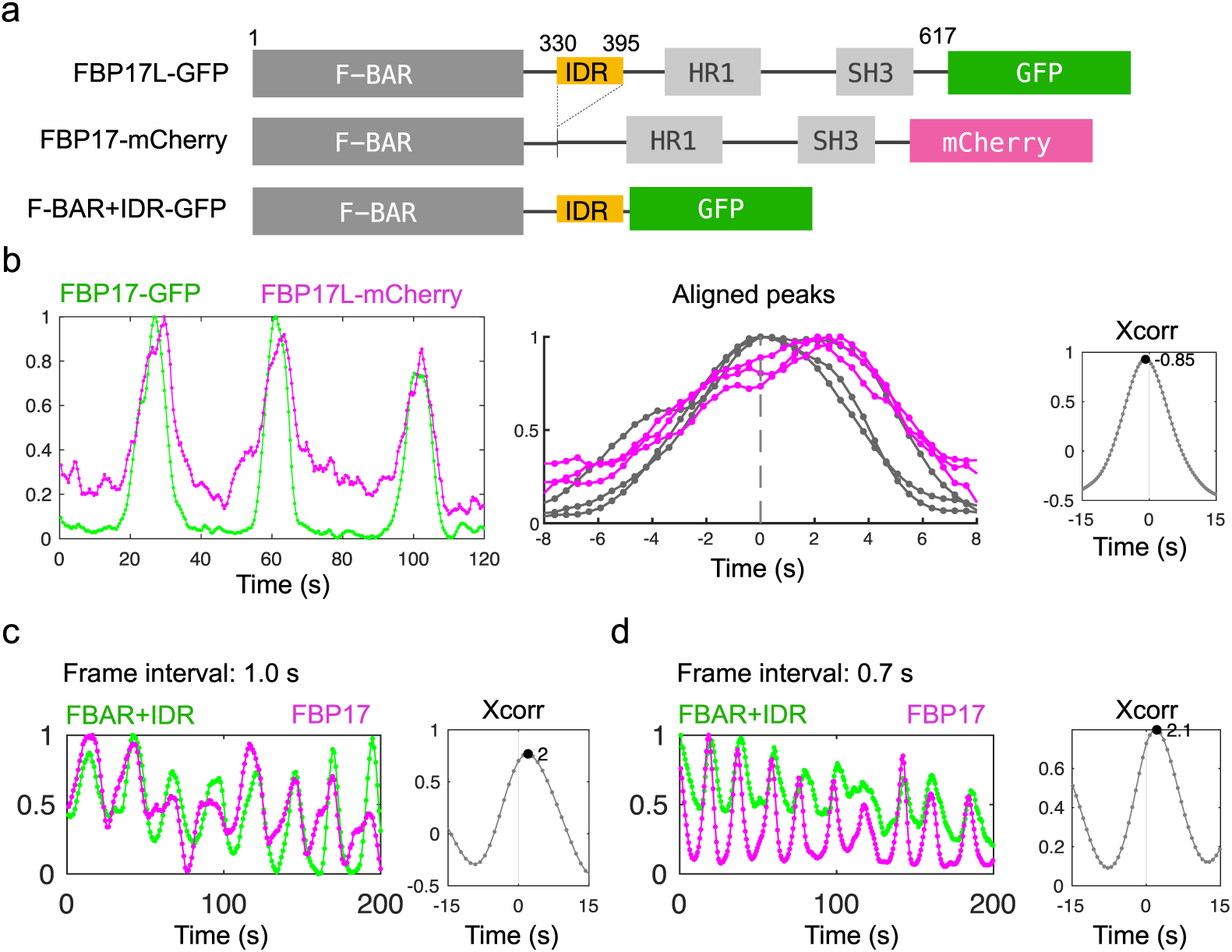
Phase difference of FBP17 isoforms in travelling waves. **a**. Diagram of domain structures of full-length FBP17 (FBP17L-GFP), short isoform (FBP17-mCherry), and F-BAR+IDR^1-395^-GFP used in this figure. The intrinsic disordered region (IDR) is shown in yellow. **b**. Intensity profile of waves of co-expressed FBP17-GFP and FBP17L-mCherry (**left**), aligned peaks of GFP channel (**Bottom**), and cross-correlation of GFP relative to mCherry. **c,d**. The phase lag of F-BAR+IDR^1-395^-GFP relative to FBP17-mCherry imaged at time interval of 1.0 s (**c**) and 0.7 s (**d**). **Left**: Intensity profile. **Right:** Cross-correlation of GFP relative to mCherry.

**Supplementary Figure 3.**
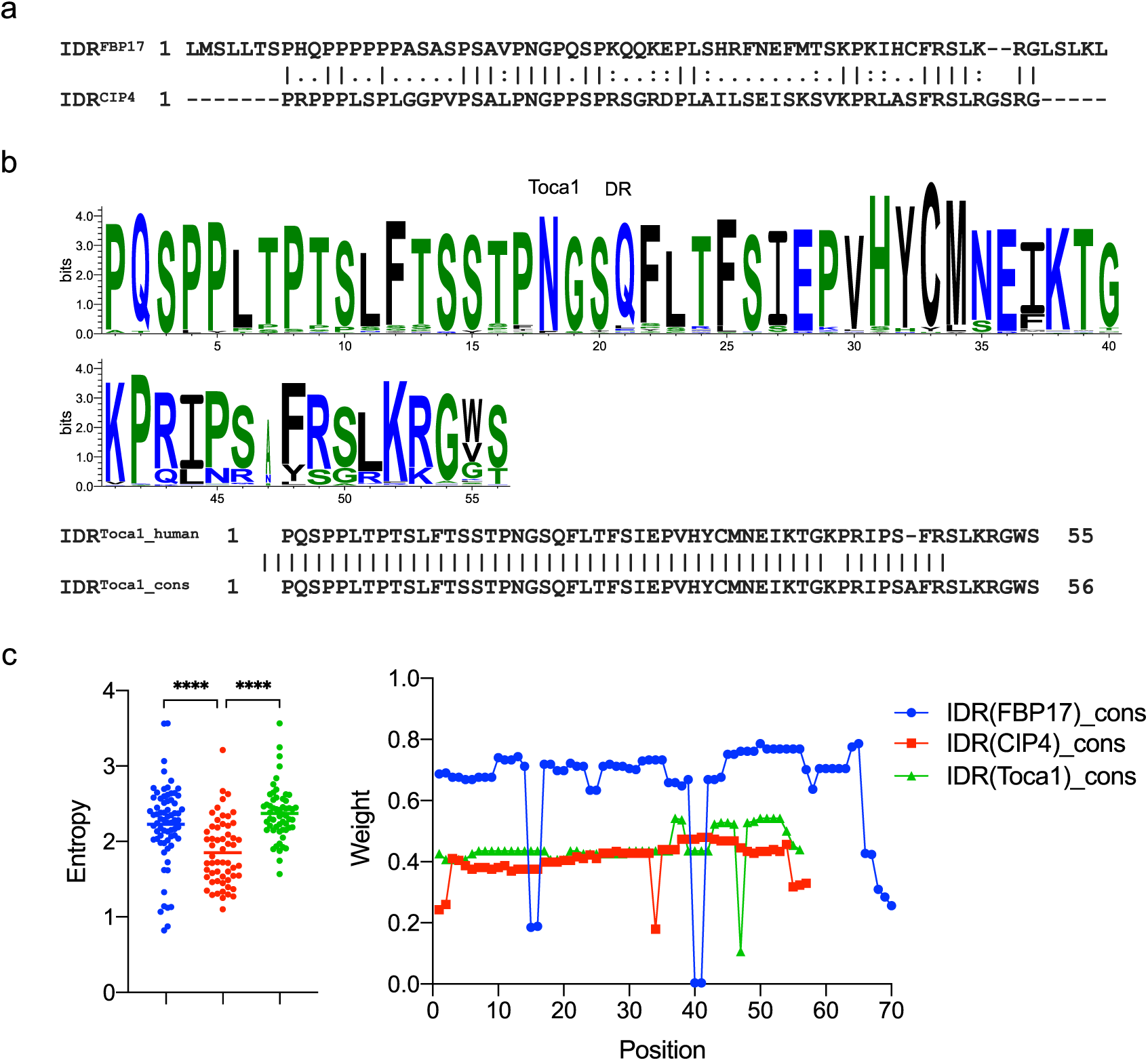
Differences between IDRs from FBP17 homologs. **a**. Sequence alignment of IDRs of human FBP17 and CIP4. **b. Top:** WebLogo of the IDR of Toca1 from all 299 species that have it. **Bottom**: Sequence alignment of the IDR from human FBP17 and the consensus sequence from WebLogo3. **c**. Sequence conservation (entropy) and splicing frequency (weight) of IDRs from FBP17, CIP4 and Toca1. Mann–Whitney U test: **** denotes P<0.0001.

